# Combining semi-automated image analysis techniques with machine learning algorithms to accelerate large scale genetic studies

**DOI:** 10.1101/152702

**Authors:** Jonathan A. Atkinson, Guillaume Lobet, Manuel Noll, Patrick E. Meyer, Marcus Griffiths, Darren M. Wells

**Affiliations:** Centre for Plant Integrative Biology, School of Biosciences, University of Nottingham, United Kingdom; Agrosphere, IBG3, Forschungszentrum Jülich, Jülich, Germany; Earth and Life Institute, Université catholique de Louvain, Louvain-la-Neuve, Belgium; InBios, Université de Liège, Liège, Belgium

**Keywords:** Root, plant phenotyping, machine learning, qtl analysis

## Abstract

**Background:** Genetic analyses of plant root system development require large datasets of extracted architectural traits. To quantify such traits from images of root systems, researchers often have to choose between automated tools (that are prone to error and extract only a limited number of architectural traits) or semi-automated ones (that are highly time consuming).

**Findings:** We trained a Random Forest algorithm to infer architectural traits from automatically-extracted image descriptors. The training was performed on a subset of the dataset, then applied to its entirety. This strategy allowed us to (i) decrease the image analysis time by 73% and (ii) extract meaningful architectural traits based on image descriptors. We also show that these traits are sufficient to identify Quantitative Trait Loci that had previously been discovered using a semi-automated method.

**Conclusions:** We have shown that combining semi-automated image analysis with machine learning algorithms has the power to increase the throughput in large scale root studies. We expect that such an approach will enable the quantification of more complex root systems for genetic studies. We also believe that our approach could be extended to other area of plant phenotyping.

## Findings

### Background

Plant root systems have many physiological roles, including the acquisition of water and nutrients, making them of critical importance for yield establishment in crops. The improvement of root architectural traits will thus be crucial in delivering the yield improvement required to ensure future global food security [1, 2]. Unfortunately, root systems are difficult to analyse and quantify: they are intrinsically complex due to their highly branched tree structure [3], and their growth in an opaque medium (soil) makes them difficult to observe.

For many years, root researchers have used specific experimental setups to observe and quantify root system architecture. Among these, the “pouch system” is widely used by the community to acquire large number of images of root systems [4–6]. In this approach, plants are grown on the surface of paper allowing the root system to be imaged. The analysis of the resulting root images can be performed either using semi-automated [7, 8] or fully-automated root image analysis software [9,10]. Semi-automated tools require input and validation by an expert user to faithfully extract the geometry of the root system. However, such user interaction is time consuming, which can strongly hinder the application of such approaches to large datasets (such as those required for quantitative genetic studies). Fully automated software tools are faster, but the extracted descriptors are prone to unexpected errors and the quantified traits are usually less informative [3]. This has led to image analysis being described as a new “bottleneck” in plant phenotyping [11].

Machine learning (an emerging multidisciplinary field of computer science, statistics, artificial intelligence, and information theory) encompasses a range of techniques for the automatic production of analytical models and has been attracting the interest of the plant science community in recent years. Machine learning is breaking new ground in plant science via the automation of procedures and experiments that previously required manual curation. These automated workflows are catalysing the development of new data driven plant science [12]; including remote sensing [13], species identification [14], and phenotyping [15–17]. Recently, a new approach utilising machine learning algorithms has been proposed for the identification of root system architectural traits. A Random Forest model was trained on corresponding ground-truth and image descriptors. The resulting trained model was used to analyse a new set of simulated images and was shown to be much more accurate than the direct image descriptors [3].

Here, we have evaluated this technique using a similar approach with experimental images, and assessed its application to a large scale genetic study. Our rationale was twofold. Firstly, we can reasonably expect a certain level of homogeneity within datasets coming from a single genetic screening as root systems from a given species share common attributes. Secondly, semi-automated root image analysis tools can be used to extract the ground-truth on a subset of images. Such ground-truths can then be used to train a machine learning algorithm that can then be used to analyse the remaining images in the dataset.

We show that such an approach can (i) yield better results than fully automated software analysis, (ii) is time-efficient compared to performing a semi-automated analysis on the whole dataset and (iii) is able to correctly identify previously found quantitative trait loci (QTL) for root traits.

### Overview of the analysis workflow

The dataset was divided in two (Fig. 1A): a training dataset *D_train_* of variable size (between 100 and 900 images out of 969, see below) and a test dataset *D_test_* of 1645 images. The test dataset consisted of images of 94 members of the winter wheat Savannah x Rialto doubled haploid mapping population, previously utilised in [5]. For each dataset, we extracted the ground-truth *T_train_, T_test_* using the semi-automated root image analysis tool RootNav [7] and a set of image descriptors *I_train_, I_test_* using a fully automated analysis pipeline, RIA-J [3] (Fig. 1B). We used the extracted data (*I_train_, T_train_*) to train a Random Forest model **M** : *I → T*, to predict the different ground-truth based on the image descriptors [3] (Fig. 1C,F). The trained Random Forest model **M** was then applied on the image descriptors *I_test_* from the test dataset *D_test_*, to predict the different ground-truth *T_test_* (named Random Forest estimators, Fig. 1D). The accuracy of both the image descriptors and the Random Forest estimators were then compared to the ground-truth acquired with RootNav.

**Figure 1:**
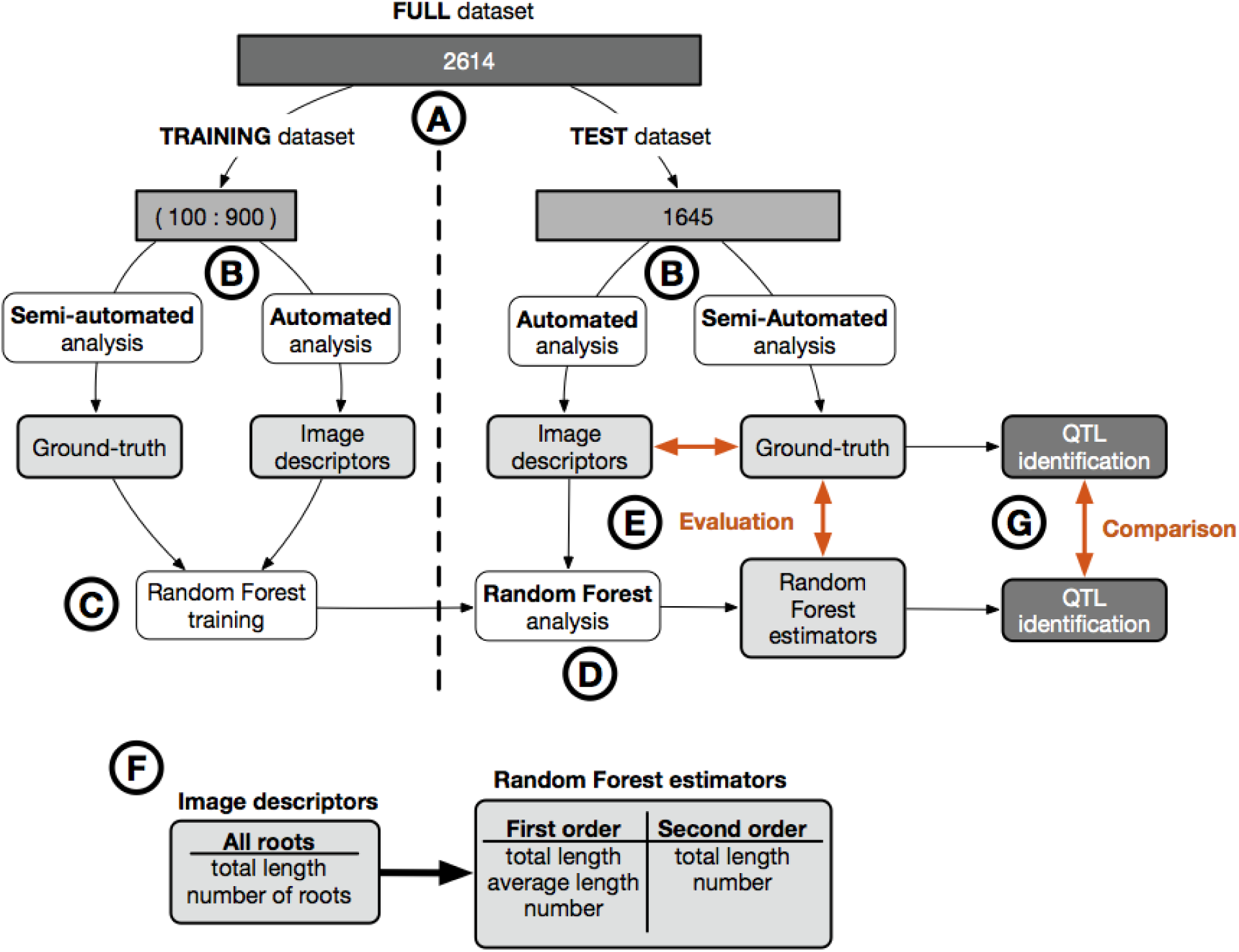
Overview of the analysis pipeline used in this study. **A.** We divided the full dataset (2614 images) into two: a training set (100 to 900 images) and a test set (1645 images). **B.** For each dataset, all the images were analysed using a semi-automated root image analysis tool (RootNav) to extract the ground-truth, as well as with a fully automated root image analysis tools (RIA-J), to extract image descriptors (see text for details). **C.** We trained a Random Forest model on the image descriptors and the ground-truth from the training dataset. **D.** We applied the Random Forest model on the image descriptors from the test dataset. **E.** We compared the image descriptors and the Random Forest estimators from the test dataset with their corresponding ground-truth. **F.** Comparison of biologically-relevant metrics extracted with the automated analysis and the Random Forest analysis. **G.** QTL were identified and compared using both Random Forest estimators and the ground-truth data.

One of the aims of our analysis was to assess the minimal size required for a training dataset. Therefore, we used different numbers of images for training: 100, 200, 300, 400, 500, 600, 700, 800 and 900. For each set, we randomly selected the images out of the 969 images that comprised the test dataset, then repeated the training/accuracy procedure described above. To account for the fact that the images were randomly selected, for each test size, we repeated the procedure 10 times.

For each test dataset size, we used the Random Forest estimators to detect QTL regions associated with the different traits quantified (Fig. 1G). The identified QTL regions were then compared to those previously identified using RootNav, as well as those identified using the direct image descriptors.

### Random Forest estimators have a greater accuracy and greater biological relevance than image descriptors

It has been previously shown that Random Forest estimators are better at predicting the ground-truth values of various root system metrics compared to direct image descriptors [3]. However, such evaluation used only simulated images, rather than a “real” experimental dataset.

Here we show that such an approach can also be used with experimental data yielding better results than the direct image descriptors (Fig. 2). We also show that, as expected, increasing the size of the training dataset increases the accuracy of the estimated metrics. For our data, we observe a strong increase in accuracy up to a dataset size of 500 test images, after which the improvement becomes marginal. Our approach also allows for the prediction of new metrics, not obtained using the direct image descriptors. For instance, the direct descriptors do not differentiate between the different root orders, whereas the Random Forest model does.

**Figure 2:**
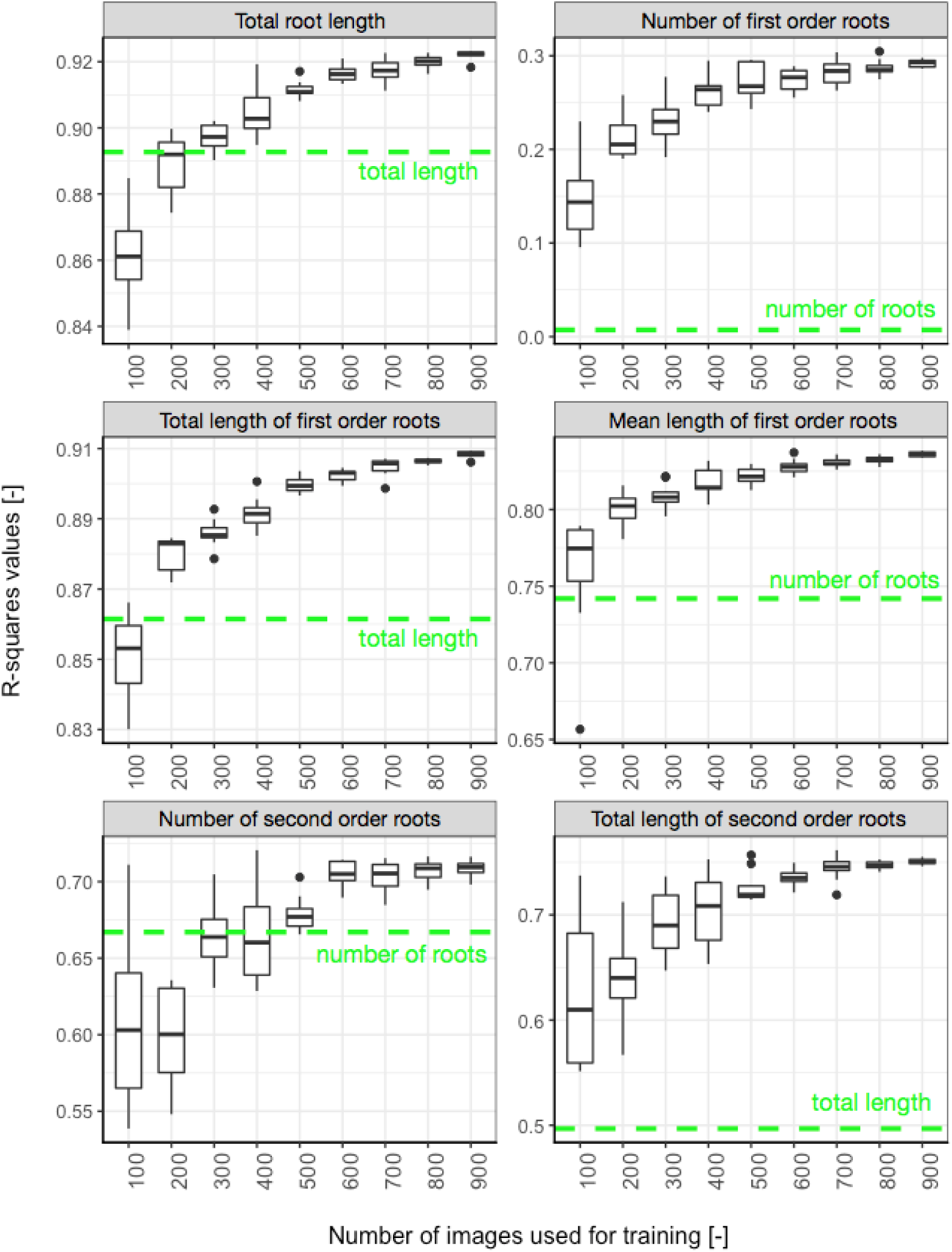
Accuracy of the Random Forest estimators. The r-squared values of the linear regression between the Random Forest estimators and the ground-truths were computed for each size and repetition of test datasets. The dotted line represents the r-squared value between the most closely related image descriptors and the ground-truth.

We observed a diminution of the range of predicted value as the number of test images increases. This may be the result of a greater accuracy of the prediction, but may also be due to the fact that the same images are randomly selected for each repetition. As the number of test images increases, we expect the number of identical images across repetitions to increase as well (the total number of test images being 969).

### Random Forest estimators identify the correct QTLs

Plant phenotyping studies often use mapping populations to dissect the genetic architecture of complex traits by identifying regions of chromosomal DNA that correlate with phenotypic variation termed quantitative trait loci (QTL). The images in our test dataset were used in such a study to identify several QTL for root traits in wheat seedlings [5]. In addition to testing the accuracy of the Random Forest approach in estimating root system parameters, we wanted to know if these parameters could be used reliably for the identification of QTL. Since QTL identification had already been performed on our test dataset, we could directly assess the performance of our new pipeline against the original approach by using the same QTL detection technique on both the direct image descriptors and the traits derived from the random forest models.

The Random Forest models, trained on different numbers of images (100:900), were used on the image descriptors from the test dataset to predict nine estimators datasets (named EST-100 to EST-900) for use in the QTL analysis (see Table 1). This was done to assess the minimum size for the training dataset required for reliable QTL detection, which may be lower than that required to accurately predict the trait values themselves. The R package R/qtl [18] was used for QTL detection on the image descriptor dataset and the nine Random Forest predicted datasets [5]. Identified QTL were then directly compared to those found in this paper.

**Table 1:** Results from the QTL comparison for the different estimator datasets: Green is a correct identification compared to results obtained using the RootNav pipeline, Red is a miss, yellow is a false positive and grey is not comparable. Numbers represent the significant LOD (logarithm of odds) score for each detected QTL generated by R/qtl. Chr: chromosome, GT: ground truth, ID: image descriptors, EST-100:900: random forest estimators derived from 100 – 900 images.

|  | Trait | ID | EST 100 | EST 200 | EST 300 | EST 400 | EST 500 | EST 600 | EST 700 | EST 800 | EST 900 | GT |
| --- | --- | --- | --- | --- | --- | --- | --- | --- | --- | --- | --- | --- |
| 4D | Width | 2.5 |  |  | 2.7 |  |  |  |  |  |  |  |
|  | W/D | 2.71 | 2.6 | 2.2 | 2.6 | 2.2 | 2.2 | 2.5 | 2.3 | 2.9 | 2.5 | 2.7 |
| 6D | Seminal number |  |  |  | 3.6 | 3.1 | 4.7 | 3.3 | 3.8 | 3.3 | 3.1 |  |
|  | Total root length | 17 | 13.6 | 15.0 | 14.4 | 15.2 | 14.2 | 16.0 | 14.7 | 16.3 | 15.3 | 24 |
|  | Mean sem. length |  | 13.4 | 13.7 | 13.8 | 13.8 | 13.8 | 14.0 | 13.9 | 13.5 | 15.6 | 22.2 |
|  | Lateral number |  | 12.6 | 19.0 | 18.2 | 17.6 | 18.5 | 17.0 | 17.6 | 16.7 | 15.4 | 9.1 |
|  | Tot lateral length |  | 11.2 | 13.0 | 14.2 | 12.0 | 15.3 | 12.6 | 13.3 | 12.2 | 11.7 | 6.4 |
|  | Tot seminal length |  | 13.8 | 13.1 | 14.1 | 14.8 | 13.7 | 15.2 | 14.7 | 14.7 | 14.4 | 25.6 |
|  | Width | 13.5 | 11.9 | 13.0 | 14.8 | 12.9 | 12.8 | 13.1 | 12.5 | 12.5 | 12.5 | 6.4 |
|  | Depth | 13.6 | 14.3 | 14.3 | 14.1 | 14.0 | 15.6 | 15.0 | 15.2 | 15.8 | 14.8 | 22.7 |
|  | W/D |  |  |  |  |  | 2.2 |  |  | 1.9 |  |  |
| 7A | Seminal number |  |  |  |  |  |  |  |  |  |  | 2.1 |
| 7D | Lateral number |  | 4.3 | 5.5 | 5.9 | 6.6 | 5.2 | 5.0 | 5.3 | 5.0 | 4.4 | 2.4 |
|  | Seminal number |  |  |  |  | 3.8 |  | 3.4 | 3.4 | 4.0 | 4.5 |  |
|  | Tot lateral length |  | 4.4 | 4.0 | 6.0 | 4.6 | 4.9 | 4.2 | 4.9 | 4.2 | 4.0 | 2 |
|  | Tot root length | 4.1 |  | 2.7 | 2.5 | 2.9 |  | 3.1 | 2.9 | 4.7 | 3.3 | 9 |
|  | Tot seminal length |  |  |  | 2.9 | 2.7 | 2.1 | 2.8 | 3.1 | 2.8 | 3.2 | 9.7 |
|  | **SUM** | **6** | **10** | **11** | **14** | **14** | **13** | **14** | **14** | **15** | **14** | **13** |

We observed that 12/13 of the expected QTL were correctly identified using the estimators from the Random Forest models trained on 600 or more images (EST-600:EST-900). We also observed that even using the smallest training set of 100 images (EST-100), most of the QTLs were identified (10/12), with 12/13 being identified with the estimators from the model trained with 300 images (EST-300). We did not observe an increase in the logarithm of odds (LOD) score with the increase of images (Table 1).

In addition, 4 extra QTL were identified on chromosomes 4D and 6D. Two of these were identified for width and width-depth ratio from EST-300, EST-500 and EST-800 datasets (Table 1). Although in this example, these have been labelled as false positives as they were not detected in the original study, they both have related QTL co-localising in the same positions (the 4D width QTL co-localises with a W/D QTL and the 6D W/D QTL co-localises with both a width and depth QTL at the same location). Both QTL were also found using the image descriptors utilised to train the Random Forest model, possibly explaining their identification. Two extra QTL for seminal number were identified on chromosomes 6D and 7D from EST-300 to EST-900 datasets. These false positives are most likely a result of the Random Forest estimators failing to assign whole numbers for seminal root count. This adds noise to the data, masking any genetic variation for seminal number, which is limited to between 4 and 6 seminal roots in this dataset.

In the majority of cases, the identified QTL had the same confidence intervals and similar peak marker positions as previously reported for all Random Forest models. Interestingly, the 4D QTL had a very similar confidence interval (position 0.8-67.6 previously reported vs 0-67.6 here), but a different peak marker position (position 4.8 previously reported vs position 30-34 here). It was also noted that lateral root QTL found on 7D had a reduced confidence interval compared to those previously reported (positions 0-101.8 previously vs 0-62.4 here).

### Combining semi-automated analysis and machine learning techniques increase the throughput of our image analysis pipeline

Extracting meaningful information from images of root systems is a subjective, tedious and often time-consuming process. As a general rule, automated techniques can only extract a limited amount of biologically relevant metrics and are often limited to young plants. Semi-automated tools are able to extract more metrics and with a greater accuracy, but at the expense of user interaction time (which makes them unsuited for large-scale genetic studies). As a result, large genetic screens targeting root system traits often focus on a set of simple traits that can be automatically extracted.

Here we have shown that machine learning techniques can be used to automatically extract a large set of root system metrics. To train the machine learning algorithm on our dataset, we estimated that 600 root images are needed. Additional images are needed to validate the accuracy of the Machine Learning estimators (around 100). These images have to be traced with a semi-automated tool to extract the parameters in the first place. Thus, instead of tracing all the images (in our case about 2600), only a subset (700) was needed. It was previously estimated that tracing one image takes, on average, 2 minutes. In our case, the whole dataset would represent a workload of 87 hours. With the combined pipeline, the workload decreased to 23 hours (27%).

In this example, we used a published dataset, for which the ground-truth data were already available [5]. In order to easily apply this approach to future studies, we have created the R application PRIMAL (Pipeline of Root Image analysis using MAchine Learning, https://plantmodelling.github.io/primal/) (Fig. 3). We recommend the following analysis strategy:

1. Use a fully automated tool to extract image descriptors global dataset.
2. Use a semi-automated tool to extract the ground-truth for 100 random images (the GROUND-TRUTH DATASET). Remove these images from the global dataset.
3. Use PRIMAL to train the Random Forest model and analyse the data.

**Figure 3:**
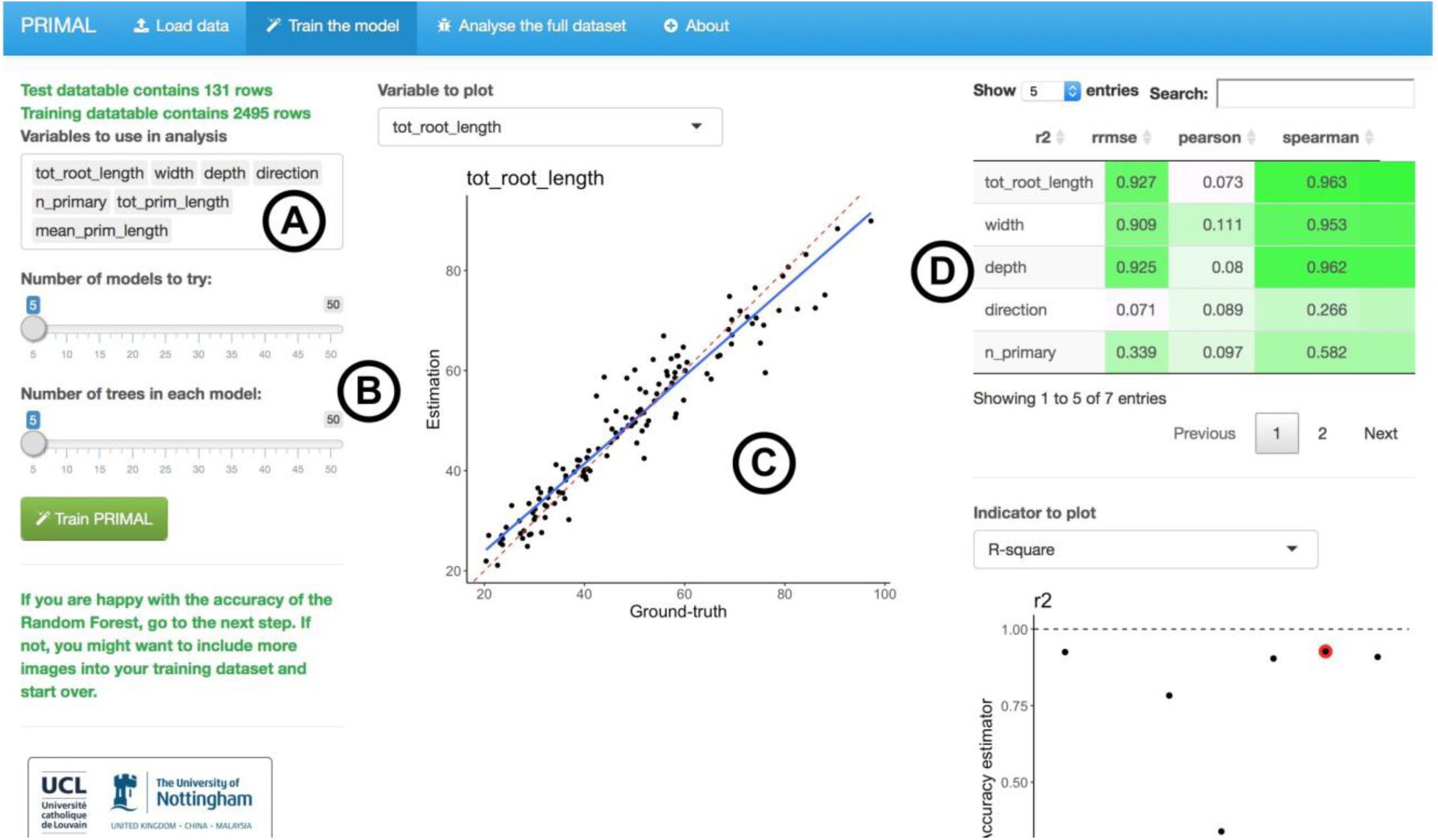
Screenshot of PRIMAL. **A.** Variable to evaluate with the Random Forest algorithm. **B.** Random Forest algorithm parameters. **C.** Visualisation of the accuracy of the Random Forest estimators. **D.** Accuracy metrics for the different descriptors.

A detailed version of this protocol is available on protocols.io: dx.doi.org/10.17504/protocols.io.h7bb9in

It should be noted that the prediction accuracy of the Random Forest estimation is highly dependent on the homogeneity of the data. For example, a Random Forest model trained on maize root systems will most likely fail when applied to wheat. However, for large scale genetic studies, where only one species is used in the analysis, this should not be an issue. The accuracy of the Random Forest estimators is also function of the variability of the direct descriptors in the dataset. Using a large set of descriptors, that better discriminate the different images, might help increase the accuracy of the Random Forest descriptors.

## Conclusions

Genetic studies on root architecture require large annotated datasets of biologically relevant traits. Automated analysis tools can be used to extract descriptors from large libraries of root images. Unfortunately, these descriptors are prone to error and their biologically relevancy is not always clear. Alternatively, semi-automated tools enable the retrieval of more precise architectural traits but, due to the requirement for skilled user inputs, they are often unsuitable for large datasets.

Here, we used a Random Forest model to predict architectural traits based on automatically-extracted image descriptors. The model was trained on a subset of the whole dataset that had been previously analysed using a semi-automated tool. This strategy allowed us to (i) decrease the time required for the analysis by 73% (compared to the semi-automated analysis of the whole dataset) and (ii) accurately predict meaningful architectural traits.

In order to make our pipeline available to the community, we have created an application available at the following address: https://plantmodelling.github.io/primal/.

## Methods

A detailed version of the protocol described here is available at Protocols.io: https://dx.doi.org/10.17504/protocols.io.h7bb9in

## Availability of supporting source code and requirements

- **Project name:** PRIMAL, Pipeline of Root Image analysis using MAchine Learning
- **Project home page:** https://plantmodelling.github.io/primal/
- **Operating system(s):** Platform independent
- **Programming language:** R
- **Other requirements:** -
- **License:** GPL

## Declarations

### List of abbreviations

QTL: Quantitative Trait Locus

### Competing interests

The author(s) declare that they have no competing interests

### Funding

MN is grateful to the F.R.S.-FNRS for a doctoral grant (1.A.320.16F).

DMW, JAA, and MG wish to acknowledge the support of the Biological and Biotechnology Science Research Council (BBSRC) for responsive mode and CISB awards to the Centre for Plant Integrative Biology and the European Research Council (ERC) for FUTUREROOTS project funding (FP7-294729).

### Authors’ contributions

- **Conceptualization**: GL, DMW, JAA
- **Formal Analysis**: GL, JAA, MN, PEM
- **Resources**: JAA, MG, MN, PEM
- **Writing – Original Draft**: GL, JAA
- **Writing – Review & Editing**: GL, JAA, MG, DMW, MN
- **Visualization**: GL, JAA

## References

1. Den Herder G, Van Isterdael G, Beeckman T, De Smet I. The roots of a new green revolution. Trends Plant Sci. 2010;15:600–7.

2. Lynch JP. Roots of the Second Green Revolution. Aust. J. Bot. CSIRO PUBLISHING; 2007;55:493–512.

3. Lobet G, Koevoets IT, Noll M, Tocquin P, Meyer PE, Pagès L, et al. Using a structural root system model to evaluate and improve the accuracy of root image analysis pipelines. Front. Plant Sci. [Internet]. Frontiers; 2017 [cited 2017 Mar 16];8. Available from: http://journal.frontiersin.org/article/10.3389/fpls.2017.00447/abstract

4. Hund A, Trachsel S, Stamp P. Growth of axile and lateral roots of maize: I development of a phenotying platform. Plant Soil. Kluwer Academic Publishers; 2009;325:335–49.

5. Atkinson JA, Wingen LU, Griffiths M, Pound MP, Gaju O, Foulkes MJ, et al. Phenotyping pipeline reveals major seedling root growth QTL in hexaploid wheat. J. Exp. Bot. Soc Experiment Biol; 2015;66:2283–92.

6. Gioia T, Galinski A, Lenz H, Müller C, Lentz J, Heinz K, et al. *GrowScreen*-*PaGe*, a non-invasive, high-throughput phenotyping system based on germination paper to quantify crop phenotypic diversity and plasticity of root traits under varying nutrient supply. Funct. Plant Biol. [Internet]. CSIRO PUBLISHING; 2016 [cited 2016 Nov 17]; Available from: http://www.publish.csiro.au.sci-hub.cc/fp/FP16128

7. Pound MP, French AP, Atkinson J, Wells DM, Bennett MJ, Pridmore T. RootNav: Navigating images of complex root architectures. 2013;162:1802–14.

8. Lobet G, Pagès L, Draye X. A novel image-analysis toolbox enabling quantitative analysis of root system architecture. Plant Physiol. American Society of Plant Biologists; 2011;157:29–39.

9. Leitner D, Felderer B, Vontobel P, Schnepf A. Recovering root system traits using image analysis exemplified by two-dimensional neutron radiography images of lupine. Plant Physiol. 2014;164:24–35.

10. Cai J, Zeng Z, Connor JN, Huang CY, Melino V, Kumar P, et al. RootGraph: a graphic optimization tool for automated image analysis of plant roots. J. Exp. Bot. Soc Experiment Biol; 2015;66:6551–62.

11. Minervini M, Scharr H, Tsaftaris SA. Image Analysis: The New Bottleneck in Plant Phenotyping [Applications Corner]. IEEE Signal Process. Mag. 2015;32:126–31.

12. Ma C, Zhang HH, Wang X. Machine learning for Big Data analytics in plants. Trends Plant Sci. 2014;19:798–808.

13. Ali I, Greifeneder F, Stamenkovic J, Neumann M, Notarnicola C. Review of Machine Learning Approaches for Biomass and Soil Moisture Retrievals from Remote Sensing Data. Remote Sensing. Multidisciplinary Digital Publishing Institute; 2015;7:16398–421.

14. Babatunde OH, Armstrong L, Leng J, Diepeveen D. A computer-based vision systems for automatic identification of plant species using kNN and genetic PCA. Jaina [Internet]. 2015;6. Available from: http://journal.magisz.org/index.php/jai/article/view/164

15. Minervini M, Abdelsamea MM, Tsaftaris SA. Image-based plant phenotyping with incremental learning and active contours. Ecol. Inform. 2014/9;23:35–48.

16. Singh A, Ganapathysubramanian B, Singh AK, Sarkar S. Machine Learning for High-Throughput Stress Phenotyping in Plants. Trends Plant Sci. Elsevier Ltd; 2015;1–15.

17. Weiss U, Biber P, Laible S, Bohlmann K, Zell A. Plant Species Classification Using a 3D LIDAR Sensor and Machine Learning. 2010 Ninth International Conference on Machine Learning and Applications. 2010. p. 339–45.

18. Broman KW, Wu H, Sen S, Churchill GA. R/qtl: QTL mapping in experimental crosses. Bioinformatics. 2003;19:889–90.

16. Ma C, Zhang H, Wang X. Machine learning for Big Data analytics in plants. Trends Plant Sci. Vol 19(12); 2014; 798–808

17. Ali I, Greifeneder F, Stamenkovic J, Neumann M, Notarnicola C. Review of Machine Learning Approaches for Biomass and Soil Moisture Retrievals from Remote Sensing Data. Remote Sensing. 2015.

18. Babatunde O, Armstrong L, Diepeveen D, Leng J. A survey of computer-based vision systems for automatic identification of plant species. Journal of Agricultural Informatics. 2015 Vol. 6, No. 1

